# Dysregulation of “Don’t Eat Me” Signaling–Related Genes in Sepsis: A Targeted Transcriptomic Analysis

**DOI:** 10.64898/2026.02.25.707968

**Authors:** Yuhai Dang, Jinliang Kong

## Abstract

Sepsis remains a life-threatening condition with limited therapeutic options targeting immune dysregulation. The CD47-SIRPα “don’t eat me” signaling axis, well characterized in tumor immune evasion, has not been systematically investigated in the context of sepsis. In this study, we performed a targeted transcriptomic analysis of phagocytosis- and “don’t eat me” –related genes using the GSE228541 dataset (14 sepsis patients, 15 healthy controls). We identified 8 significantly differentially expressed genes within the curated gene panel. Key changes included downregulation of CD47 (logFC = −0.88, FDR = 5.6 × 10^−4^) and marked upregulation of PRTN3 (logFC = 2.68, FDR = 6.1 × 10^−4^). Gene Ontology (GO) enrichment demonstrated prominent alterations in pathways including negative regulation of phagocytosis (GO:0050765, FDR = 7.6 × 10^−22^), endocytosis, and inflammatory responses. Co-expression network analysis identified SNX3, DYSF, and PLSCR1 as hub genes within this regulatory module. Immune infiltration analysis showed increased M1 macrophage polarization and neutrophil activation in sepsis. Using LASSO regression, we constructed a 6-gene diagnostic signature (PLSCR1, SNX3, DYSF, PRTN3, CSK, CD47) that discriminated sepsis from controls with good performance (AUC = 0.933 in the test subset). Downregulation of CD47 suggests impaired “self” recognition, which may contribute to aberrant phagocytosis during sepsis. Elevated PRTN3 is consistent with neutrophil activation and extracellular trap formation, linking innate immune activation to tissue injury. This targeted transcriptomic analysis reveals coordinated transcriptional reprogramming of phagocytosis-regulatory genes in sepsis and supports the CD47-SIRPα axis as a candidate therapeutic target for further investigation.

## Introduction

Sepsis is defined as life-threatening organ dysfunction caused by a dysregulated host response to infection, and remains a leading cause of mortality in intensive care units worldwide despite extensive research and therapeutic advances [1]. The pathophysiology of sepsis involves complex interactions between excessive inflammation, counter-regulatory immunosuppression, coagulopathy, and endothelial injury, yet the molecular mechanisms underlying these processes remain incompletely understood.

The “don’t eat me” signal, primarily mediated by the **CD47-SIRPα axis**, serves as a fundamental mechanism protecting healthy cells from phagocytic clearance [2]. CD47, a ubiquitously expressed transmembrane protein, interacts with signal regulatory protein α (SIRPα) on macrophages and dendritic cells to deliver an inhibitory signal that suppresses phagocytosis. This pathway has been extensively studied in cancer, where tumor cells frequently upregulate CD47 to evade immune surveillance, and CD47 blockade has emerged as a promising immunotherapeutic strategy [3].

However, the expression and functional role of “don’t eat me” signaling in acute inflammatory disorders such as sepsis remain **poorly defined**. Given the profound immune dysregulation observed in sepsis—including cytopenias, macrophage reprogramming, and altered phagocytic activity—we hypothesized that genes involved in “don’t eat me” signaling and phagocytosis regulation may be transcriptionally dysregulated. Emerging evidence suggests that immune checkpoint pathways may contribute to sepsis-associated immunosuppression [4], and neutrophil extracellular traps (NETs) are strongly implicated in sepsis-induced organ dysfunction [5,6].

In this study, we performed a **targeted transcriptomic analysis** of genes associated with “don’t eat me” signaling and phagocytosis regulation using the publicly available GSE228541 dataset [7]. We characterized differential expression patterns, performed functional enrichment and co-expression network analyses, evaluated immune infiltration profiles, and explored a potential transcriptional signature for sepsis discrimination with validated visualizations of key analytical results.

## Methods

### Data Acquisition and Processing

The gene expression dataset GSE228541 was obtained from the Gene Expression Omnibus (GEO) database [7]. This dataset includes RNA-sequencing profiles from 14 sepsis patients and 15 healthy controls. Raw read counts were retrieved and processed for downstream analysis.

A **curated gene panel** was constructed based on biological relevance to “don’t eat me” signaling, phagocytosis regulation, and membrane-mediated recognition processes. The panel included 12 genes: CD47, PLSCR1, SNX3, TLR2, DYSF, TGFB1, HMGB1, PRTN3, FCGR2B, CSK, ATG5, and ATG3.

### Differential Expression Analysis

Expression counts were normalized and analyzed using the **limma-voom framework** appropriate for RNA-seq data [8]. Lowly expressed genes were filtered prior to analysis (mean count < 1 in fewer than 2 samples). Linear modeling and empirical Bayes moderation were used to identify genes differentially expressed between sepsis and control groups. P-values were adjusted for multiple testing using the Benjamini–Hochberg false discovery rate (FDR) procedure. Genes with |logFC| > 0.5 and FDR < 0.05 were considered significantly differentially expressed.

### Functional Enrichment Analysis

Gene Ontology (GO) Biological Process enrichment analysis was performed using the clusterProfiler package [9]. Enriched terms were identified using hypergeometric testing with FDR correction. Terms with adjusted p < 0.05 were considered statistically significant. A gene-pathway network was constructed to visualize the relationships between dysregulated genes and enriched biological pathways.

### Co-Expression Network Analysis

A **gene co-expression network** was constructed based on pairwise Pearson expression correlations among the curated genes. Edges were retained for pairs with |correlation coefficient| > 0.6. Network topology parameters, including node degree centrality, were calculated to identify highly connected hub genes. Nodes were colored by expression change (upregulated/downregulated) and edges by correlation direction (positive/negative) for visualization.

### Immune Infiltration Analysis

Relative abundances of immune cell subsets were estimated using**single-sample gene set enrichment analysis (ssGSEA)** based on established immune cell signatures. Enrichment scores for major immune cell types were calculated and compared between sepsis and control groups to characterize immune infiltration patterns.

### Diagnostic Signature Development

Least Absolute Shrinkage and Selection Operator (LASSO) regression was applied using the glmnet package [10] to identify a compact gene signature from the curated panel for discriminating sepsis from controls. Ten-fold cross-validation was used to select the optimal penalty parameter (λ) by minimizing binomial deviance. Random forest analysis was performed with 500 trees to evaluate feature importance. The dataset was randomly split into training (70%) and testing (30%) sets. Model performance was assessed using receiver operating characteristic (ROC) curve analysis.

### Statistical Analysis

All statistical analyses were performed in R version 4.3. Continuous variables were compared using the Mann– Whitney U test. A two-sided p < 0.05 was considered statistically significant unless otherwise specified.

## Results

### Differential Expression of “Don’t Eat Me” –Related Genes in Sepsis

Within the curated 12-gene panel, **8 genes were significantly differentially expressed** between sepsis patients and healthy controls (FDR < 0.05; Table 1, Figure 1). CD47, the core “don’t eat me” signal molecule, was significantly downregulated in sepsis (logFC = −0.88, FDR = 5.6 × 10^−4^), suggesting reduced expression of the “self” recognition signal. PRTN3, which encodes neutrophil proteinase 3, was strongly upregulated (logFC = 2.68, FDR = 6.1 × 10^−4^), consistent with neutrophil activation and NET formation.

**Table 1.** Differential expression of “don’t eat me” –related genes in sepsis patients versus healthy controls. Abbreviations: logFC, log2(fold change); AveExpr, average expression level; P.Value, raw p-value; adj.P.Val, adjusted p-value (FDR, Benjamini–Hochberg); Up, upregulated in sepsis; Down, downregulated in sepsis; Not Sig, not significant.

| Gene | logFC | AveExpr | P.Value | adj.P.Val | Regulation |
| --- | --- | --- | --- | --- | --- |
| SNX3 | 1.48 | 11.90 | $2.74 \times 10^{-6}$ | $3.29 \times 10^{-5}$ | Up |
| DYSF | 1.64 | 10.90 | $8.03 \times 10^{-6}$ | $4.82 \times 10^{-5}$ | Up |
| PLSCR1 | 1.63 | 11.01 | $1.74 \times 10^{-5}$ | $7.00 \times 10^{-5}$ | Up |
| CD47 | −0.88 | 10.89 | $1.88 \times 10^{-4}$ | $5.65 \times 10^{-4}$ | Down |
| PRTN3 | 2.68 | 4.60 | $2.55 \times 10^{-4}$ | $6.11 \times 10^{-4}$ | Up |
| CSK | −0.72 | 10.13 | $1.20 \times 10^{-3}$ | $2.41 \times 10^{-3}$ | Down |
| TGFB1 | −0.65 | 7.33 | $1.82 \times 10^{-3}$ | $3.12 \times 10^{-3}$ | Down |
| ATG3 | 0.68 | 10.27 | $6.12 \times 10^{-3}$ | $9.17 \times 10^{-3}$ | Up |
| TLR2 | 0.49 | 11.55 | $8.43 \times 10^{-2}$ | $1.12 \times 10^{-1}$ | Not Sig |
| ATG5 | −0.48 | 7.69 | $1.44 \times 10^{-1}$ | $1.73 \times 10^{-1}$ | Not Sig |
| HMGB1 | -0.15 | 11.39 | $3.34 \times 10^{-1}$ | $3.64 \times 10^{-1}$ | Not Sig |
| FCGR2B | -0.01 | 9.17 | $9.78 \times 10^{-1}$ | $9.78 \times 10^{-1}$ | Not Sig |

*Statistical criteria:* $|\log FC| > 0.5$ and adj.P.Val (FDR) $< 0.05$ .

**Figure 1.**
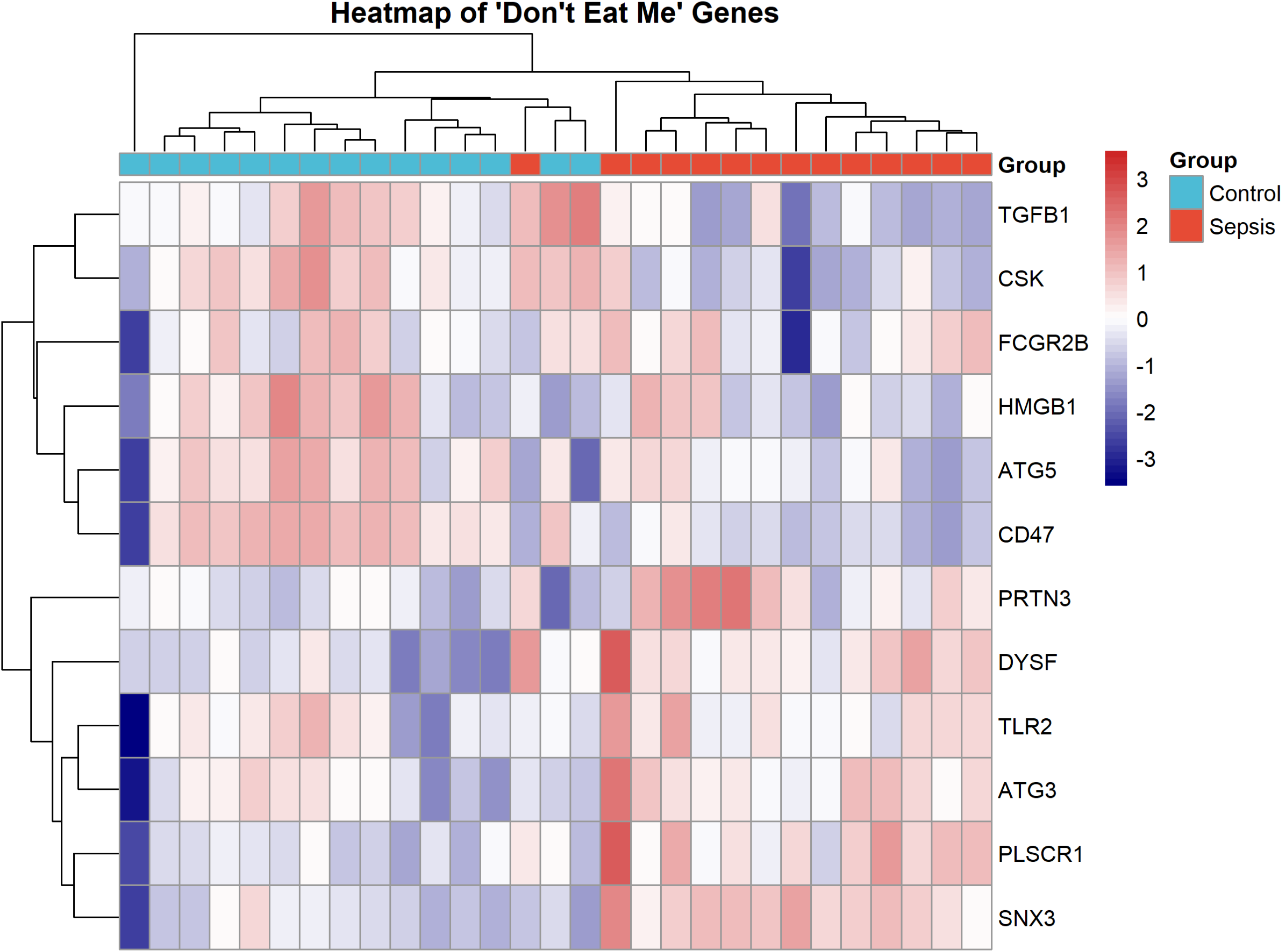
Expression patterns of “don’t eat me” genes in sepsis. (A) Volcano plot showing differential expression. Red: upregulated (FDR<0.05, logFC>0.5); Blue: downregulated (FDR<0.05, logFC<-0.5). (B) Box plots of key differentially expressed genes. (C) Heatmap of all 12 genes with hierarchical clustering.

Other significantly upregulated genes included **SNX3, DYSF, and PLSCR1**, which are involved in membrane trafficking, endosomal sorting, and phagocytic regulation. Significantly downregulated genes included **CSK and TGFB1**. ATG3 was modestly upregulated, while TLR2, ATG5, HMGB1, and FCGR2B did not reach statistical significance.

### Functional Enrichment Indicates Dysregulated Phagocytosis

GO enrichment analysis of the curated gene set revealed strong enrichment for phagocytosis-related biological processes (Table 2, Figure 2B-C). The most significantly enriched term was **negative regulation of phagocytosis** (GO:0050765, FDR = 7.6 × 10^−22^). Additional enriched terms included negative regulation of endocytosis (GO:0045806, FDR = 1.6 × 10^−17^), regulation of phagocytosis (GO:0050764, FDR = 2.8 × 10^−16^), and phagocytosis (GO:0006909, FDR = 2.1 × 10^−13^). We also observed enrichment for “regulation of Fc receptor mediated stimulatory signaling pathway” (GO:0060368, FDR = 1.1 × 10^−6^) and interleukin-6 production pathways (GO:0032635, FDR = 4.2 × 10^−3^). These results indicate that sepsis is associated with **coordinated transcriptional dysregulation of multiple phagocytosis-related pathways**.

**Table 2.**
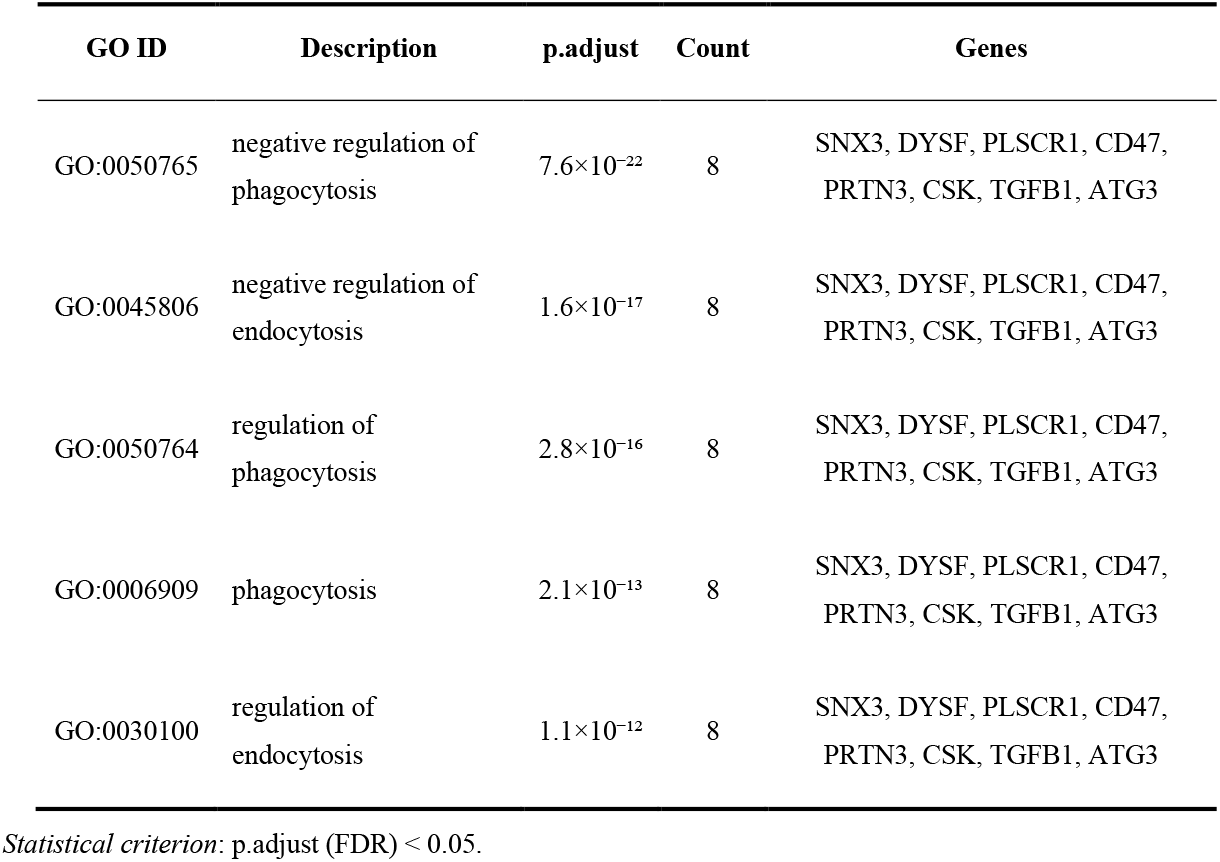
Top enriched Gene Ontology Biological Process terms. Abbreviations: GO, Gene Ontology; p.adjust, adjusted p-value (FDR); Count, number of input genes involved in the term.

| GO ID | Description | p.adjust | Count | Genes |
| --- | --- | --- | --- | --- |
| GO:0050765 | negative regulation of phagocytosis | $7.6 \times 10^{-22}$ | 8 | SNX3, DYSF, PLSCR1, CD47, PRTN3, CSK, TGFB1, ATG3 |
| GO:0045806 | negative regulation of endocytosis | $1.6 \times 10^{-17}$ | 8 | SNX3, DYSF, PLSCR1, CD47, PRTN3, CSK, TGFB1, ATG3 |
| GO:0050764 | regulation of phagocytosis | $2.8 \times 10^{-16}$ | 8 | SNX3, DYSF, PLSCR1, CD47, PRTN3, CSK, TGFB1, ATG3 |
| GO:0006909 | phagocytosis | $2.1 \times 10^{-13}$ | 8 | SNX3, DYSF, PLSCR1, CD47, PRTN3, CSK, TGFB1, ATG3 |
| GO:0030100 | regulation of endocytosis | $1.1 \times 10^{-12}$ | 8 | SNX3, DYSF, PLSCR1, CD47, PRTN3, CSK, TGFB1, ATG3 |

*Statistical criterion:* p.adjust (FDR) $< 0.05$ .

**Figure 2.**
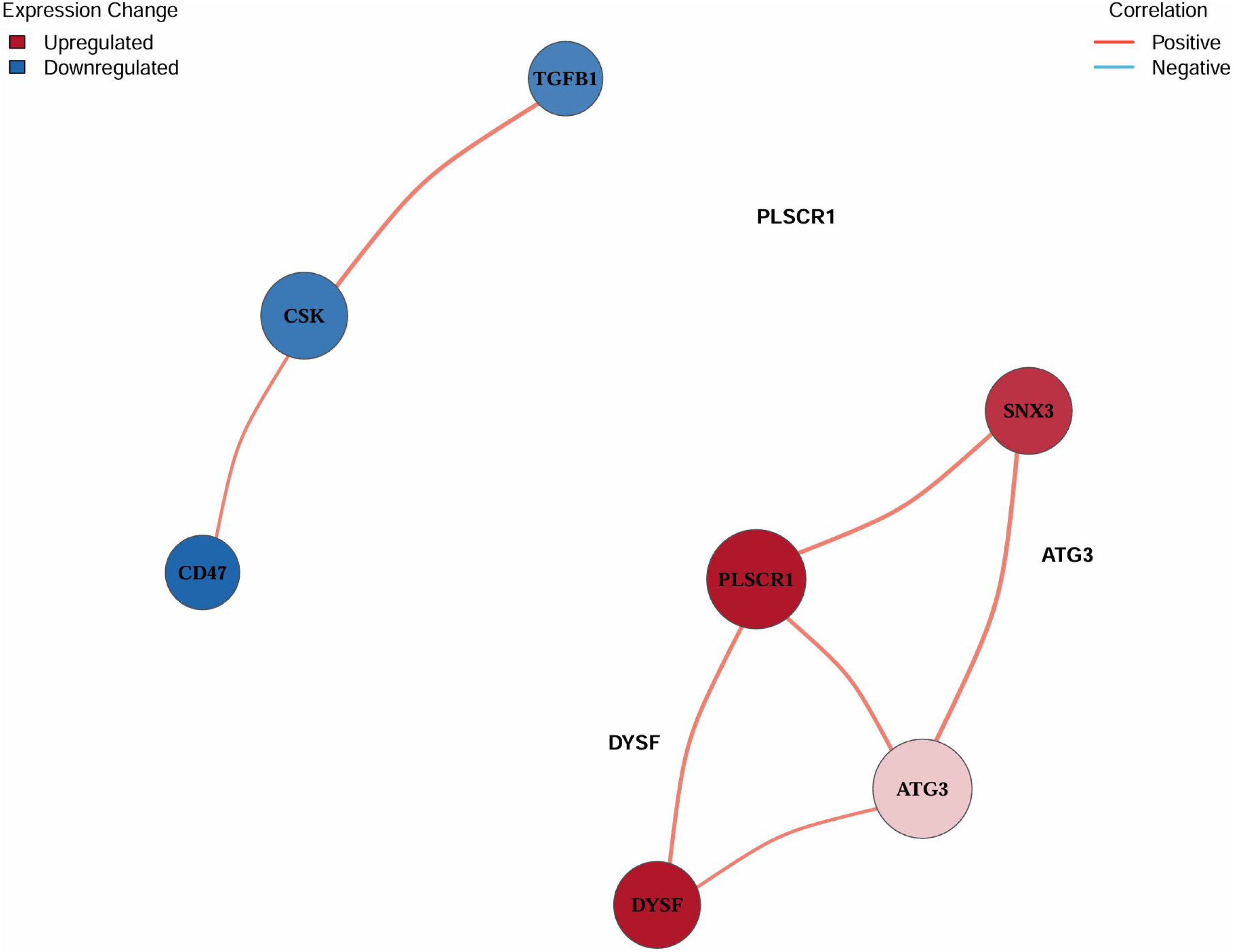
Functional analysis of differentially expressed genes. (A) Co-expression network of “don’t eat me” –related genes. Node color indicates expression change (red = upregulated in sepsis, blue = downregulated in sepsis). Edge color indicates correlation direction (positive = red, negative = blue). Nodes represent core genes in the phagocytosis regulatory module, with hub genes (SNX3, DYSF, PLSCR1) showing the highest connectivity. (B) GO Biological Process enrichment bubble plot. (C) GO enrichment bar plot. (D) Gene-pathway network illustrating the molecular relationships between dysregulated “don’t eat me” –related genes (purple nodes) and enriched GO Biological Process terms (yellow nodes) in sepsis. Edges represent the association of individual genes with specific biological pathways, highlighting the central involvement of core genes (CD47, PLSCR1, PRTN3, SNX3, DYSF) in phagocytosis, endocytosis, and inflammatory signaling pathways.

A gene-pathway network further illustrated the specific connections between dysregulated genes and enriched biological processes, with core genes such as CD47, PLSCR1, and PRTN3 mapping to key phagocytosis and inflammatory pathways (Figure 2D).

### Co-Expression Network Identifies Hub Genes

Co-expression network analysis revealed strong interconnections among the dysregulated genes (Figure 2A). Nodes were colored by expression change (upregulated/downregulated) and edges by the direction of expression correlation (positive/negative). **SNX3, DYSF, and PLSCR1** were identified as hub genes with the highest connectivity. CD47 remained connected to these core genes, supporting its role within a broader phagocytosis-regulatory module. PRTN3 showed correlations with other network components, linking neutrophil activation to membrane and repair pathways.

### Immune Infiltration Patterns in Sepsis

ssGSEA-based immune infiltration analysis revealed **elevated M1 macrophage and neutrophil signatures** in sepsis patients compared with controls, consistent with a pro-inflammatory innate immune phenotype (Figure 3). Monocyte signatures were also increased, reflecting activation of the mononuclear phagocyte system—an observation consistent with the upregulation of neutrophil-associated genes such as PRTN3 in sepsis.

**Figure 3.**
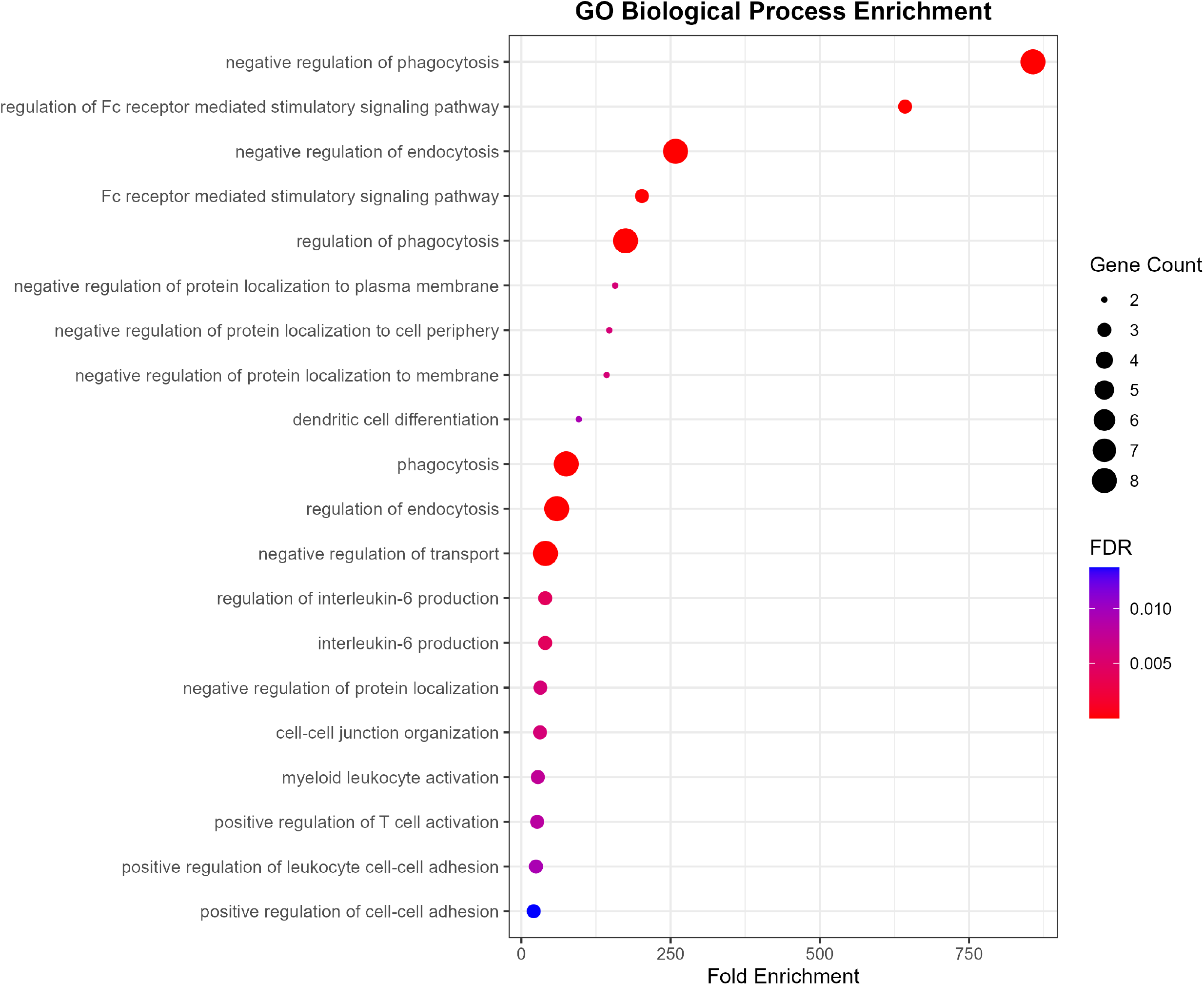
Immune infiltration patterns. Heatmap and box plots showing relative abundance of immune cell types estimated by ssGSEA. Sepsis samples exhibit elevated M1 macrophage and neutrophil infiltration.

### Diagnostic Signature Performance

LASSO regression with ten-fold cross-validation was used to select the optimal penalty parameter (λ), with the cross-validation plot showing the relationship between −log(λ) and binomial deviance (Figure 4A). This analysis identified a **6-gene signature** (PLSCR1, SNX3, DYSF, PRTN3, CSK, CD47) that distinguished sepsis from control samples, with positive coefficients for upregulated genes (PLSCR1, SNX3, DYSF, PRTN3) and negative coefficients for downregulated genes (CSK, CD47).

**Figure 4.**
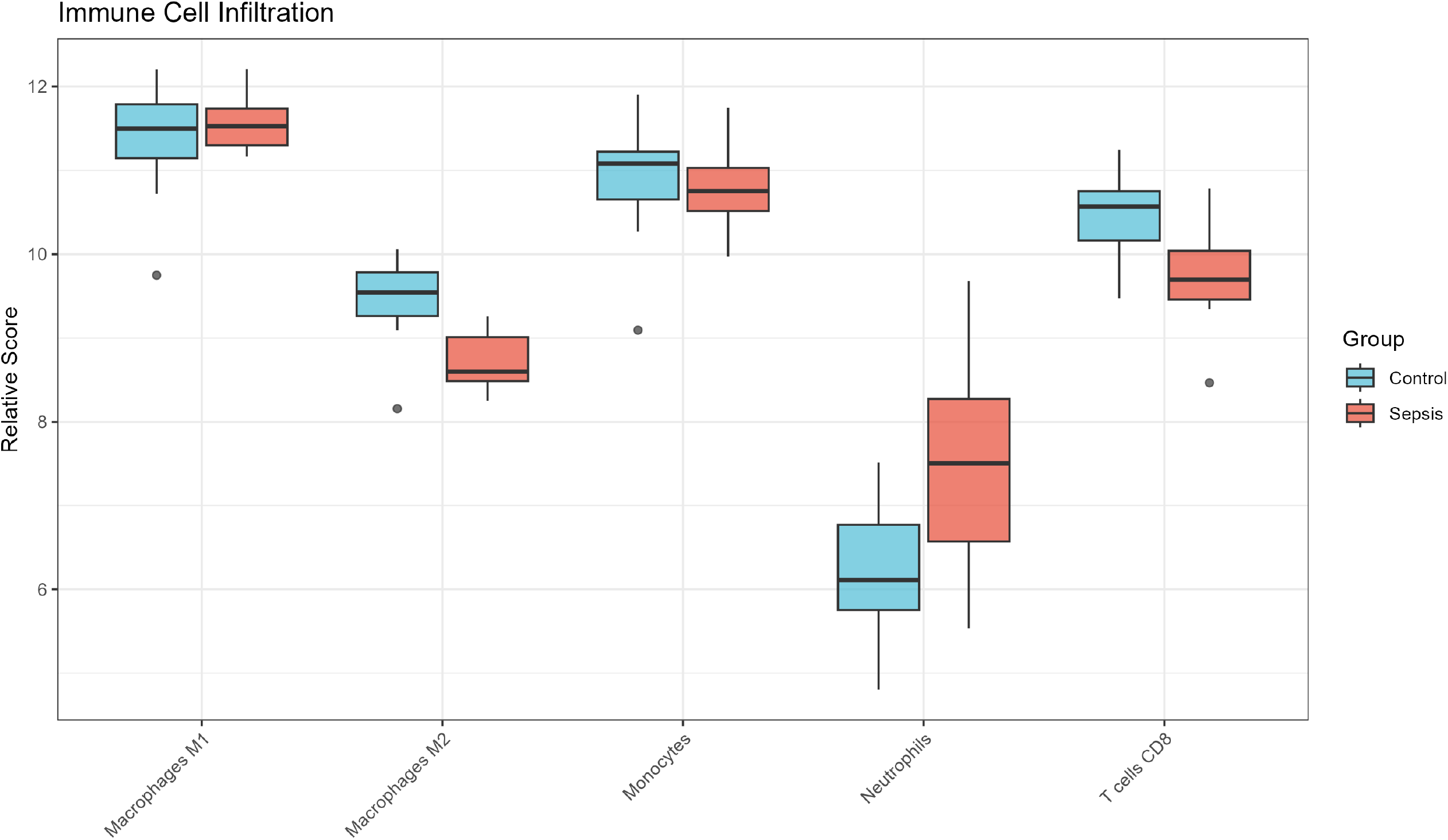
Diagnostic model performance. (A) LASSO cross-validation plot for penalty parameter (λ) selection. Ten-fold cross-validation plot showing the relationship between −log(λ) (x-axis) and binomial deviance (y-axis) for the LASSO regression model. The plot was used to select the optimal penalty parameter that minimizes deviance, guiding the identification of the 6-gene diagnostic signature for sepsis. The x-axis also displays the number of genes retained in the model at each λ value (top). (B) Random forest feature importance. (C) ROC curves for LASSO and random forest models on test set. Receiver operating characteristic (ROC) curves evaluating the discriminative performance of the LASSO 6-gene signature and random forest model in the test subset (30% of the dataset). The y-axis represents sensitivity, and the x-axis represents specificity. The LASSO model achieved an AUC of 0.933, and the random forest model achieved an AUC of 1.0, demonstrating strong performance in distinguishing sepsis patients from healthy controls.

Random forest analysis highlighted **SNX3, CD47, and DYSF** as the most influential features for classification accuracy (Table 3, Figure 4B). In the test subset, the LASSO model achieved an **AUC of 0.933**, while the random forest model reached an AUC of 1.0 (Figure 4C), supporting the potential of these transcriptional signatures as classifiers for sepsis.

**Table 3.** Feature importance from random forest analysis. Abbreviation: Mean Decrease Gini, measure of variable importance for classification performance (higher values indicate greater contribution to model accuracy).

| Gene | Mean Decrease Gini |
| --- | --- |
| SNX3 | 2.85 |
| CD47 | 2.74 |
| DYSF | 2.42 |
| PLSCR1 | 1.55 |
| PRTN3 | 1.04 |
| CSK | 0.89 |
| TGFB1 | 0.76 |
| ATG3 | 0.54 |
| ATG5 | 0.43 |
| TLR2 | 0.31 |
| HMGB1 | 0.22 |
| FCGR2B | 0.16 |

## Discussion

This study provides a **targeted transcriptomic analysis** of “don’t eat me” and phagocytosis-related genes in sepsis, with validated visualizations of key analytical results, revealing coordinated transcriptional changes in key regulatory pathways. The findings generate important hypotheses about the role of phagocytosis regulation in sepsis pathophysiology and identify potential candidate biomarkers and therapeutic targets.

The **downregulation of CD47** represents a notable observation in this cohort. In cancer, CD47 upregulation promotes immune evasion by inhibiting phagocytic clearance of tumor cells, whereas in sepsis, reduced CD47 expression may**diminish “self” protection** of host cells. This impairment in the core “don’t eat me” signal could potentially contribute to inappropriate phagocytosis of healthy host cells, which may underlie the cytopenias and endothelial damage characteristic of severe sepsis [4]. These findings align with emerging evidence linking CD47-dependent signaling to immune regulation in sepsis and support further investigation of this axis in acute inflammatory disease.

The marked **upregulation of PRTN3** in sepsis aligns with the well-established role of neutrophils and NETs in sepsis pathophysiology [5,6]. PRTN3, a key component of neutrophil azurophilic granules, contributes to NET structure and function; its upregulation thus links innate immune activation to tissue injury, coagulopathy, and organ dysfunction—hallmarks of severe sepsis. This observation reinforces the importance of neutrophil dysregulation in sepsis and identifies PRTN3 as a potential marker of neutrophil activation in this condition.

Hub genes **SNX3, DYSF, and PLSCR1** identified in the co-expression network (Figure 2A) highlight the importance of **membrane dynamics, trafficking, and repair** in phagocytic regulation during sepsis. Their coordinated dysregulation suggests widespread remodeling of cell surface and vesicular processes that govern phagocytosis and immune recognition. The gene-pathway network (Figure 2D) further contextualizes these genes within key biological processes, demonstrating their involvement in phagocytosis, endocytosis, and inflammatory signaling—all pathways central to sepsis pathophysiology.

The 6-gene transcriptional signature showed **good discriminative performance** in this cohort, with the LASSO model achieving an AUC of 0.933 and the random forest model an AUC of 1.0 in the test subset (Figure 4C). The LASSO cross-validation plot (Figure 4A) confirms the robust selection of the penalty parameter for model construction, and random forest analysis identifies the most biologically relevant features for classification. However, given the small sample size and single-dataset design, these results should be considered**preliminary and hypothesis-generating**, and validation in larger, independent cohorts is essential to confirm the signature’s diagnostic utility.

A key translational implication of these findings is that the **CD47-SIRPα signaling axis may represent a potential therapeutic target** in sepsis. Notably, the therapeutic direction for sepsis may be opposite to that in cancer: enhancing CD47-mediated “don’t eat me” signaling could potentially reduce aberrant phagocytosis of host cells and protect vulnerable tissues from immune-mediated damage, a strategy that warrants further investigation in preclinical sepsis models.

### Limitations

This study has several important limitations. First, it is a **retrospective, targeted analysis** restricted to a predefined 12-gene set, rather than an unbiased genome-wide discovery design, which may limit the identification of additional novel genes and pathways involved in sepsis-associated phagocytosis dysregulation. Second, the **sample size is small** (n = 29), with no independent external validation cohort, increasing the risk of overfitting—particularly evident in the perfect AUC of the random forest model (Figure 4C). Third, findings are limited to **transcript levels** and lack protein expression or functional validation; future studies should confirm these observations at the protein level and perform functional assays to validate the role of the CD47-SIRPα axis in sepsis. Fourth, SIRPA, the canonical receptor for CD47, was not reliably detected in this dataset, precluding analysis of the complete signaling axis. Fifth, the observational design does not establish causal relationships between gene dysregulation and sepsis pathophysiology.

Despite these limitations, this study provides a**systematic characterization of “don’t eat me” –related gene dysregulation** in sepsis, with rigorous visualization of key analytical results, and generates testable hypotheses for future mechanistic and translational studies.

## Conclusions

This targeted transcriptomic analysis demonstrates significant dysregulation of “don’t eat me” and phagocytosis-related genes in sepsis, characterized by **CD47 downregulation and PRTN3 upregulation** as key features. Co-expression network analysis identifies SNX3, DYSF, and PLSCR1 as central hub genes in a phagocytosis-regulatory network (Figure 2A), and a gene-pathway network (Figure 2D) contextualizes these genes within core biological pathways disrupted in sepsis. LASSO regression identifies a 6-gene signature (PLSCR1, SNX3, DYSF, PRTN3, CSK, CD47) that shows good discriminative ability in this cohort, with the LASSO model achieving an AUC of 0.933 in the test subset (Figure 4C) and robust penalty parameter selection via ten-fold cross-validation (Figure 4A). These results support further investigation of the CD47-SIRPα axis as a **potential therapeutic target** and the 6-gene signature as a candidate biomarker for sepsis, with validation in larger, independent cohorts a critical next step.

## Acknowledgments

The authors are grateful for the support and collaboration of The First Affiliated Hospital of Guangxi Medical University throughout this study. We extend our sincere thanks to the Department of Medical Records and the
clinical teams for their invaluable assistance in data acquisition and curation. We also acknowledge the use of public databases, including GeneCards and GEO, for providing valuable bioinformatic resources. Finally, we
deeply appreciate all the patients who participated in this research.

## Author Contributions

Conceptualization: Y.D., J.K.; Data analysis: Y.D.; Visualization: Y.D.; Writing - original draft: Y.D.; Writing - review & editing: J.K.; Supervision: J.K.; Funding acquisition: J.K.

## Ethics approval and consent to participate

Not applicable.

## Data Availability

The datasets analyzed in this study are available from the Gene Expression Omnibus (GEO) under accession number GSE228541. Processed data and analysis scripts are available from the corresponding author upon
reasonable request.

---

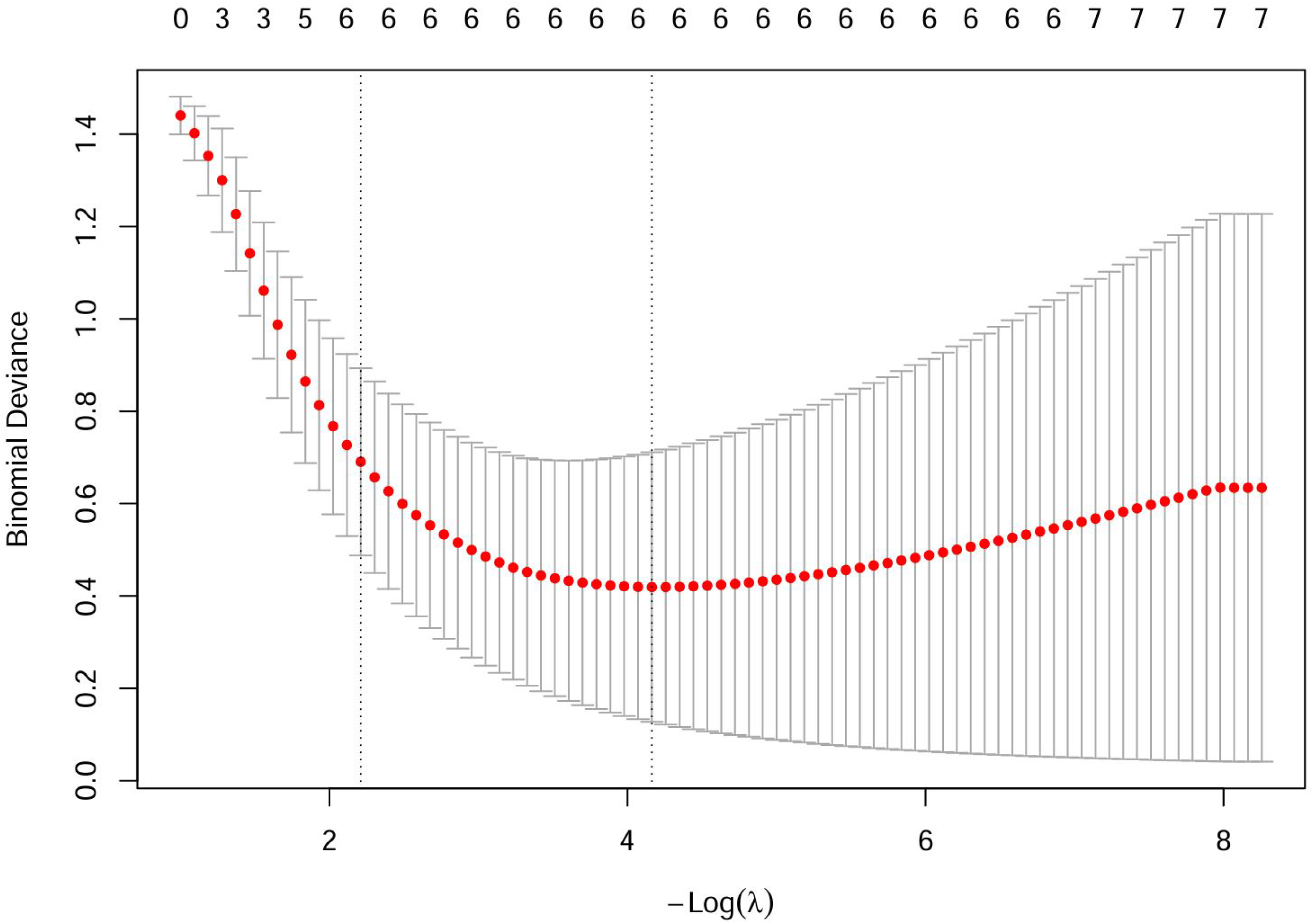

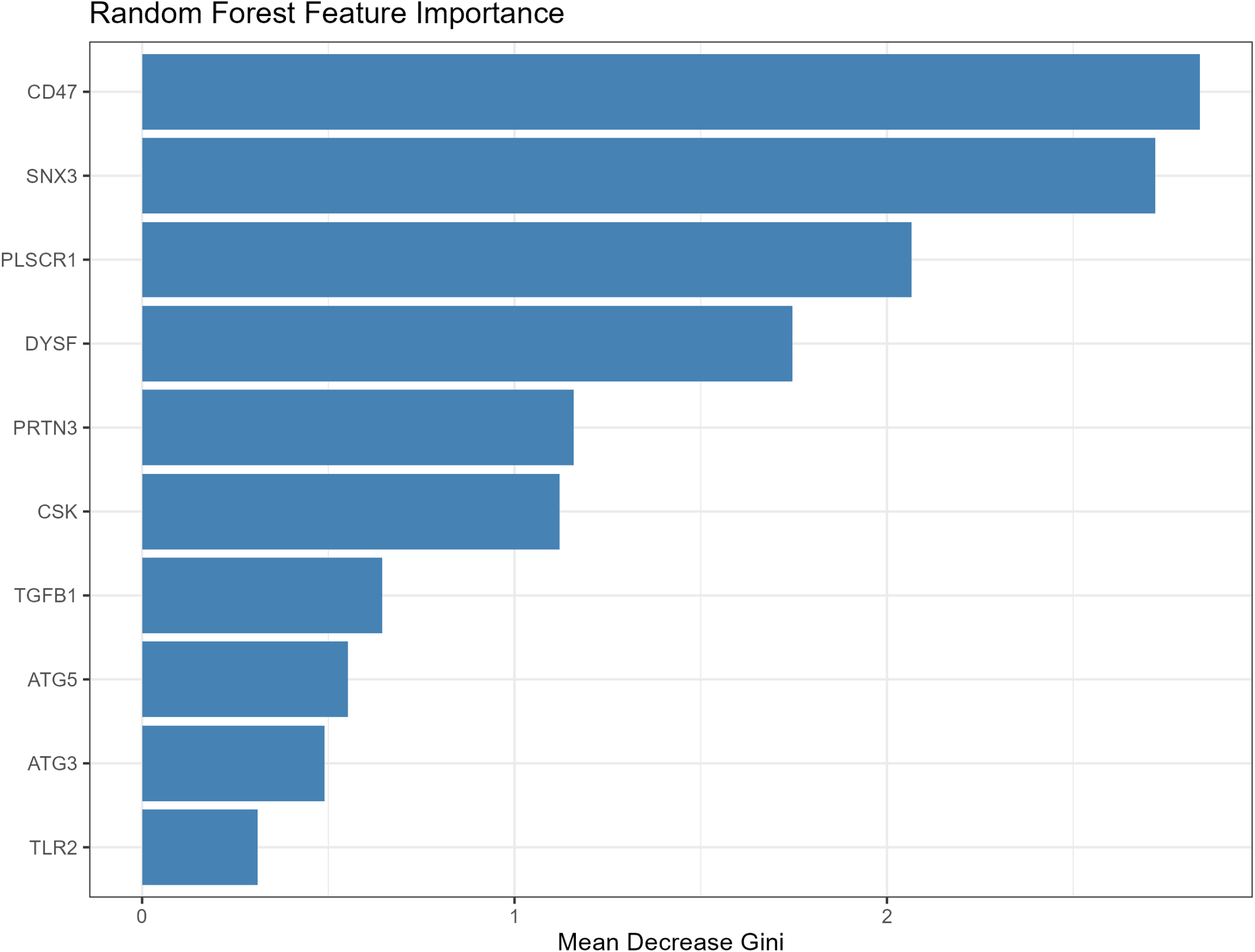

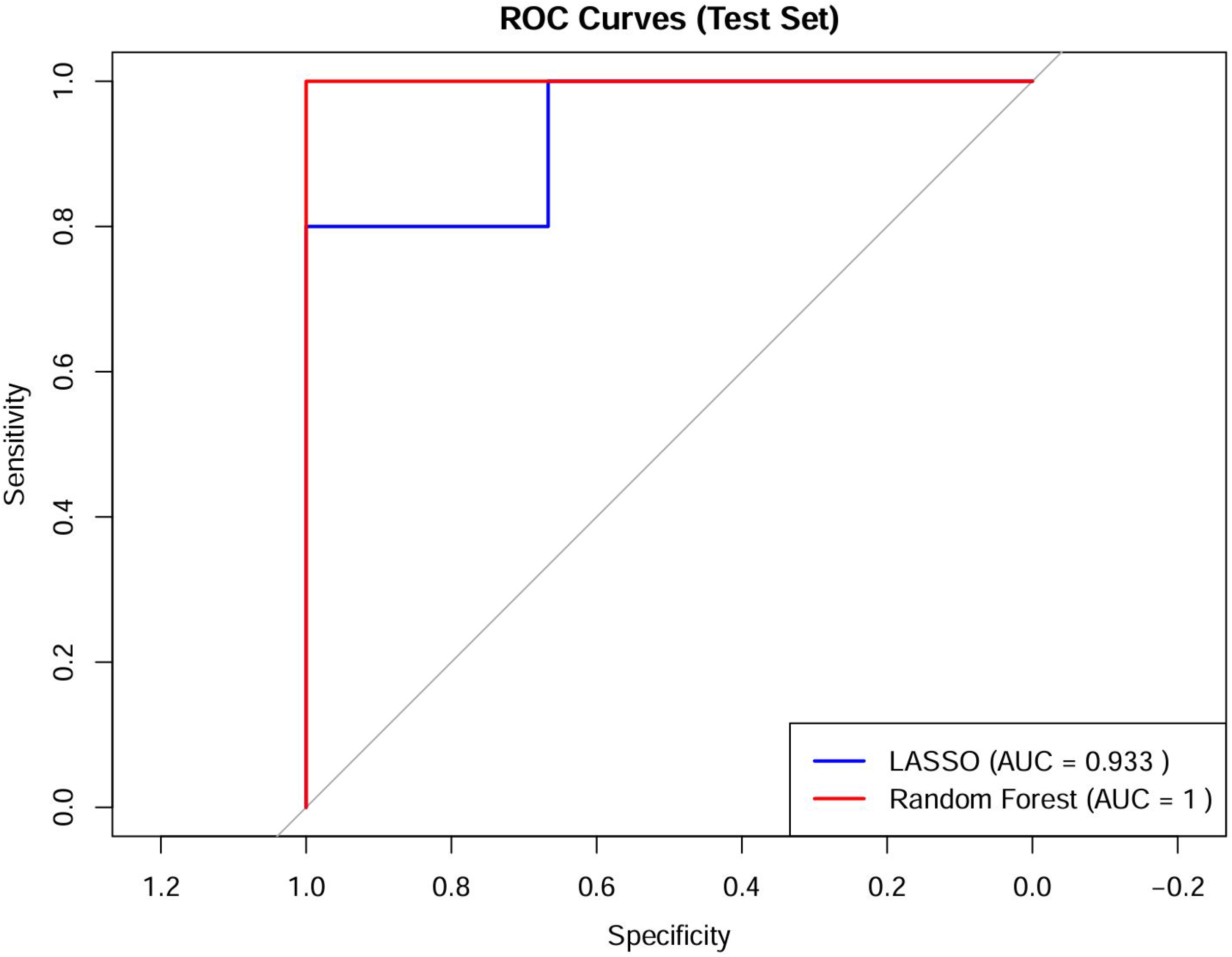

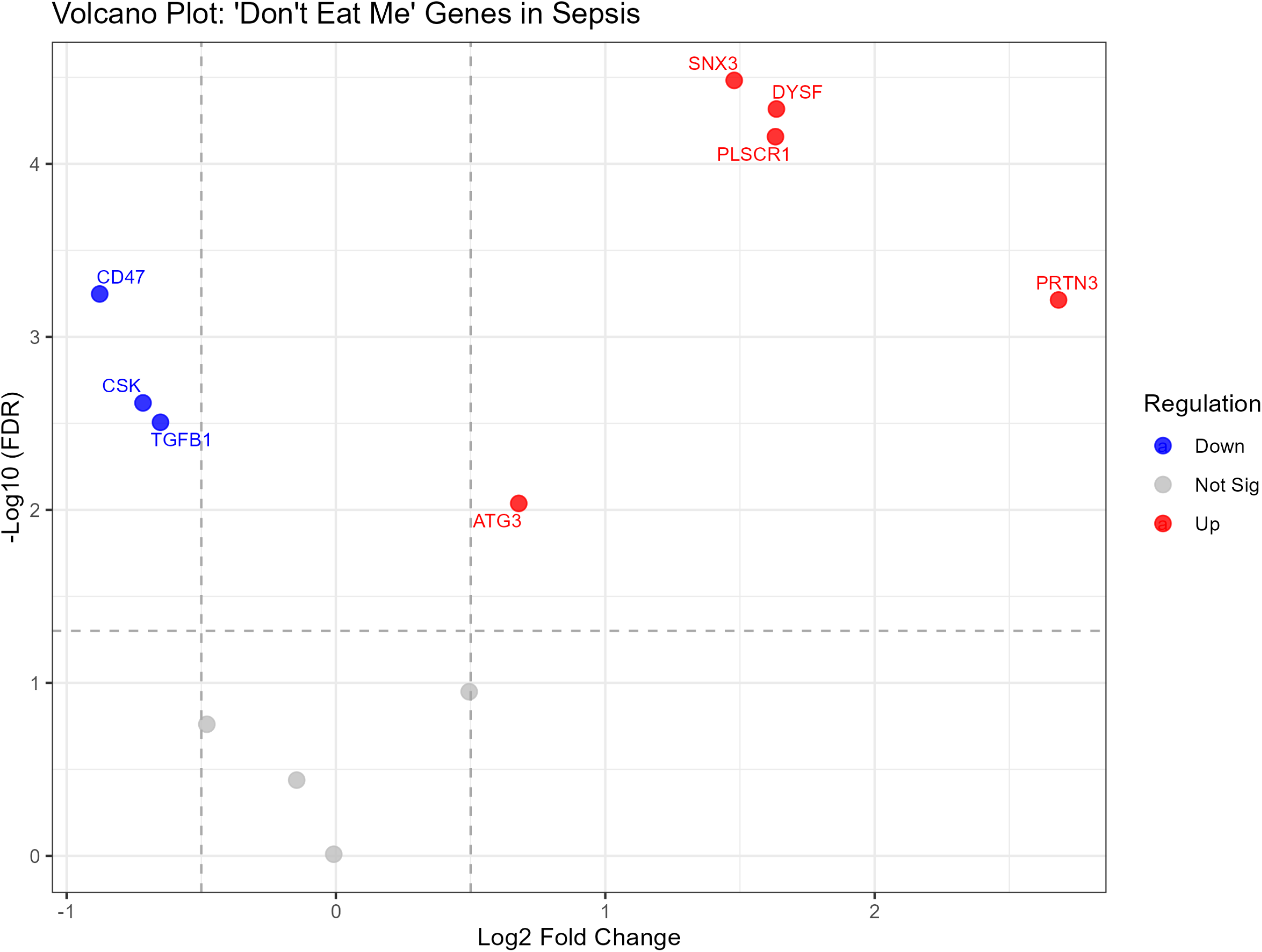

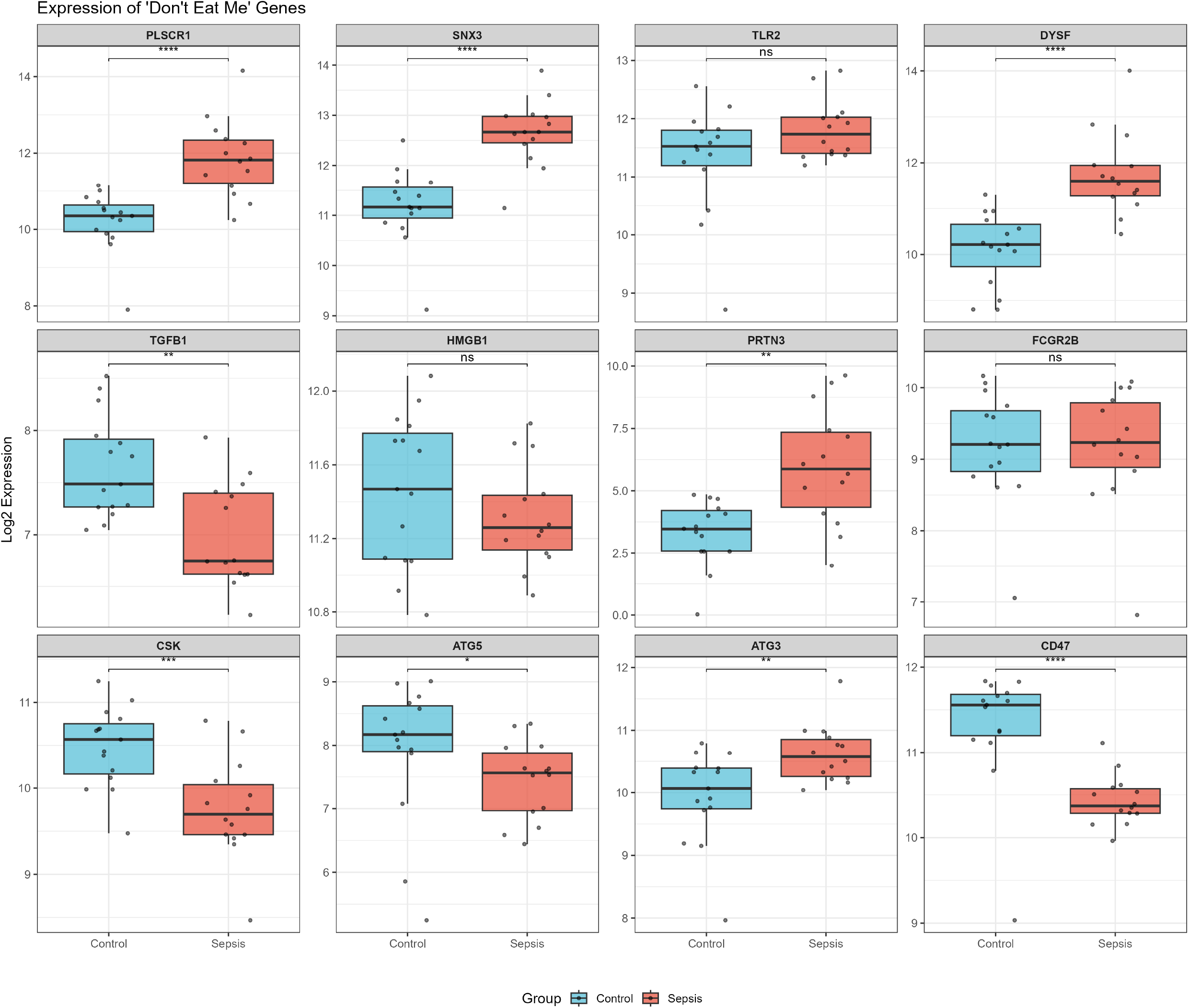

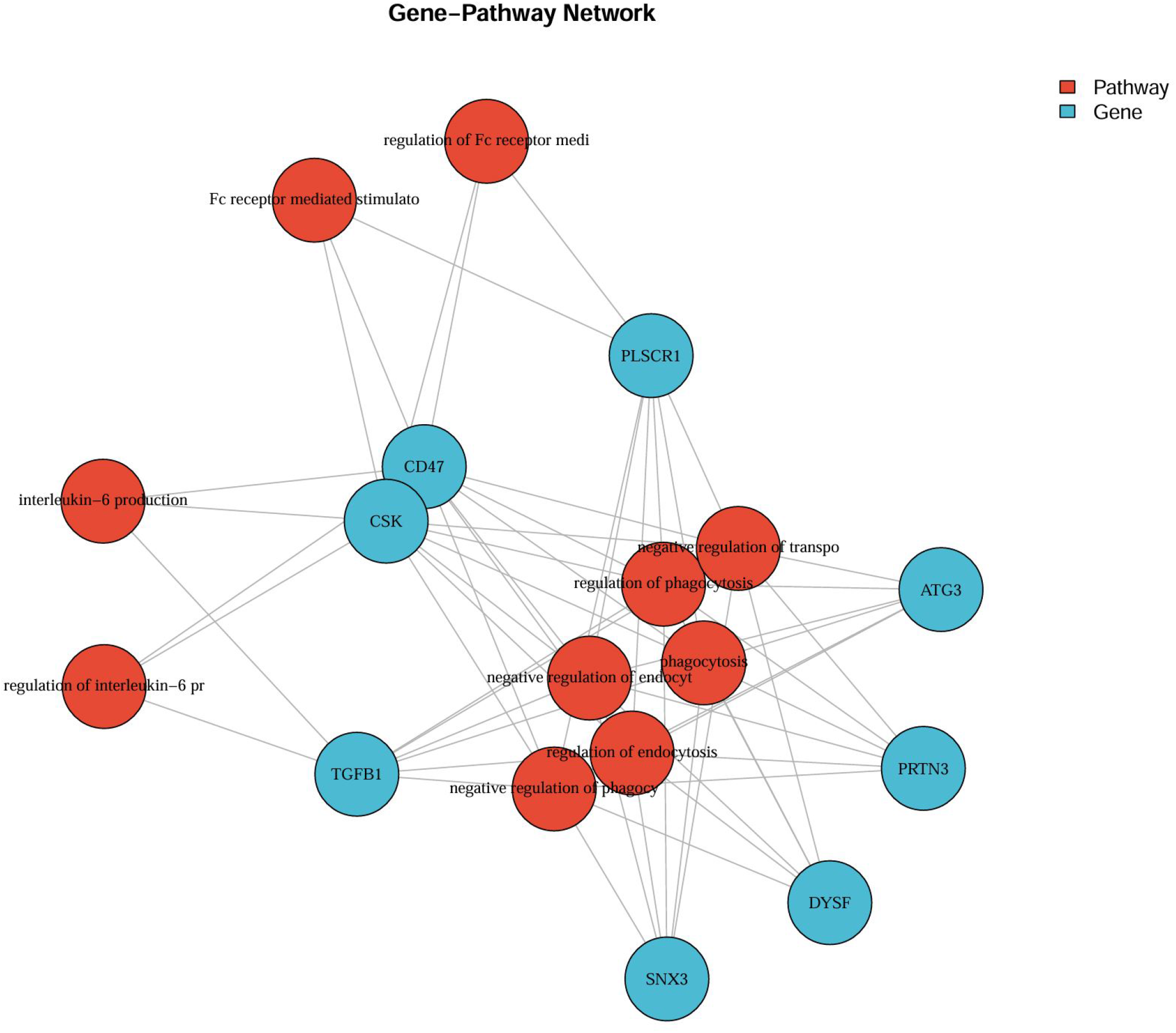

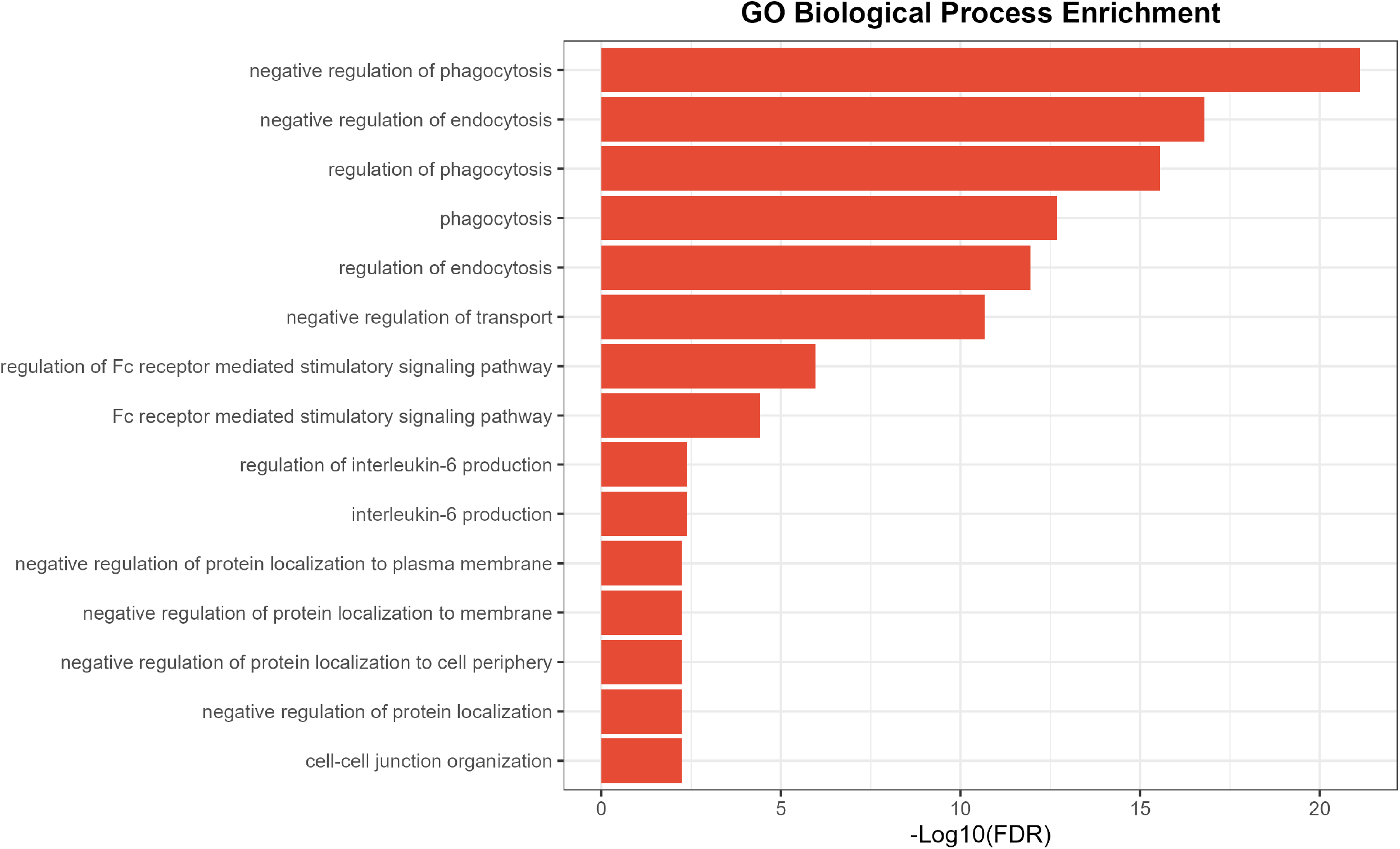

